# Modular, Vascularized Hypertrophic Cartilage Constructs for Bone Tissue Engineering Applications

**DOI:** 10.1101/2024.02.26.582166

**Authors:** Nicholas G. Schott, Gurcharan Kaur, Rhima Coleman, Jan P. Stegemann

**Author notes:** equal contribution.

## Abstract

Insufficient vascularization is a main barrier to creating engineered bone grafts for treating large and ischemic defects. Modular tissue engineering approaches have promise in this application because of the ability to combine tissue types and to localize microenvironmental cues to drive desired cell function. In direct bone formation approaches, it is challenging to maintain sustained osteogenic activity, since vasculogenic cues can inhibit tissue mineralization. This study harnessed the physiological process of endochondral ossification to create multiphase tissues that allowed concomitant mineralization and vessel formation. Mesenchymal stromal cells in pellet culture were differentiated toward a cartilage phenotype, followed by induction to chondrocyte hypertrophy. Hypertrophic pellets exhibited increased alkaline phosphatase activity, calcium deposition, and osteogenic gene expression relative to chondrogenic pellets. In addition, hypertrophic pellets secreted and sequestered angiogenic factors, and supported new blood vessel formation by co-cultured endothelial cells and undifferentiated stromal cells. Multiphase constructs created by combining hypertrophic pellets and vascularizing microtissues and maintained in unsupplemented basal culture medium were shown to support robust vascularization and sustained tissue mineralization. These results demonstrate a new *in vitro* strategy to produce multiphase engineered constructs that concomitantly support the generation of mineralize and vascularized tissue in the absence of exogenous osteogenic or vasculogenic medium supplements.

**Graphical Abstract:** 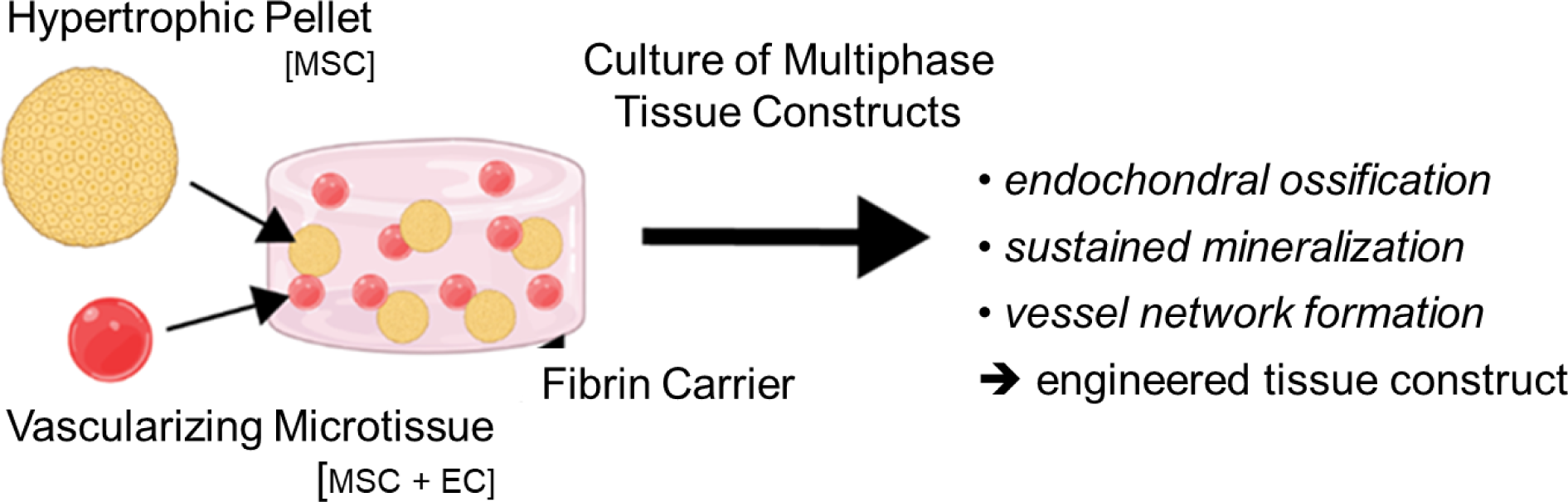

## 1. Introduction

Cell-based tissue engineering has the potential to transform the treatment of complex musculoskeletal pathologies, such as large bone defects and fracture non-unions. However, the achievement of timely and adequate vascularization of complex engineered tissues has been identified as a main challenge in achieving clinical translation [1,2]. Most approaches to prevascularizing engineered bone tissues have involved the addition of a vasculogenic phase to a preformed osteogenic construct, and subsequent culture in a ‘hybrid’ nutrient medium containing both osteoinductive components and vasculogenic factors [3–6]. However, it has been shown that key components in endothelial cell growth medium inhibit osteogenic differentiation of MSC, even when the medium is also supplemented with osteoinductive factors [7]. Furthermore, the addition of osteoinductive factors has been reported to compromise EC proliferation, migration, and the formation of vessel-like structures [8,9]. These mutually incompatible effects pose a significant challenge in efforts to couple vascular and osseous tissue development for regenerative therapies. A variety of prevascularization strategies designed to create a vascular network within engineered constructs prior to transplantation have been investigated [10], with the goal of accelerating inosculation with host vasculature. These strategies have shown that prevascularized constructs can survive *in vivo* [11,12] and may enhance neovascularization [13,14]. Adult mesenchymal stromal cells (MSC) are often used as a cell source in these applications because they are natural osteoprogenitors with a well-documented capacity to undergo controllable osteogenesis *in vitro* and support bone regeneration *in vivo* [15–18]. In addition, MSC have been widely used as supportive pericytes in vascular tissue engineering, in which they play an instrumental role in vessel formation and maturation by secreting angiogenic factors and directly interacting with endothelial cells (EC) to stabilize vasculature [19–21]. However, in bone tissue engineering applications, it has also been demonstrated that undifferentiated MSC secrete soluble factors that directly inhibit osteogenic differentiation *in vitro* [22]. Therefore, there is a need to develop strategies that allow for concomitant and sustained regeneration of both bone and vascular tissues in a unified tissue construct.

An alternate approach to creating vascularized bone constructs involves harnessing the process of endochondral ossification (EO), which is characterized by the formation of an intermediate cartilaginous template that naturally promotes vascular infiltration and matrix mineralization, which thereby generates new vascularized bone tissue [23,24]. This dynamic process involves progenitor cells undergoing chondrogenesis, leading to the production of a dense collagenous extracellular matrix rich in proteoglycans and elastin fibers [25]. Chondrocytes entrapped in this cartilaginous anlage eventually mature to hypertrophy, as marked by substantial cell enlargement and deposition of collagen type X. Hypertrophic chondrocytes secrete proangiogenic factors, including vascular endothelial growth factor (VEGF) and matrix metalloproteinases (MMPs), which promote vascular infiltration. It has been proposed that newly formed blood vessels allow the ingress of osteoclasts and osteoprogenitor cells to facilitate matrix remodeling, mineralization, and subsequent bone development [26]. More recently, it has been suggested that direct chondrocyte-to-osteoblast transdifferentiation may occur, such that hypertrophic chondrocytes directly contribute to mineralization of the formed matrix and subsequent bone development [27,28].

EO-based strategies to creating vascularized bone that have been investigated previously apply chondrogenic pellet culture of MSC to induce the formation of a cartilaginous matrix and subsequent adoption of a hypertrophic phenotype [29]. Hypertrophic pellets are then embedded within a vascularizing matrix containing EC and MSC and cultured in endothelial growth medium. This culture system is tailored to support EC viability and morphogenesis, with the goal of vascularizing the hypertrophic constructs prior to transplantation. However, studies to date have highlighted some of the challenges in applying this culture system. While the presence of pre- formed vascular networks was found to accelerate construct mineralization by four weeks of subcutaneous implantation, this effect did not correlate with enhanced vascularization *in vivo* [30]. Another study demonstrated that hypertrophic constructs transplanted with pre-formed EC networks resulted in significantly better vascularization following transplantation; however, improved bone formation was not observed, and it was postulated that components in the endothelial growth medium used to prevascularize the construct had impaired the cartilage-to-bone transition of hypertrophic chondrocytes [31]. These promising yet varied results demonstrate the need for improved culture strategies to promote concomitant tissue mineralization and new blood vessel development.

The overall objective of this study was to develop a new *in vitro* strategy to generate prevascularized bone constructs that support both mineral deposition and blood vessel development within a unified system. A modular approach emulating the EO pathway was used in which discrete populations of progenitor cell-laden microtissues were separately cultured to produce either a hypertrophic cartilage or a vascularizing tissue type. Chondrogenic pellets produced using human MSC were induced to hypertrophy to create endochondral tissues that could both mineralize and support vascularization. Hypertrophic pellets were then combined with modular microtissues containing EC and MSC to allow vessel formation. These multiphase constructs were maintained under defined culture conditions, which allowed comparison of tissue compositions and media formulations. These studies resulted in a new approach to combining modular microtissues to allow concomitant formation of vascularized and mineralized tissue in unsupplemented culture medium.

## 2. Materials and Methods

### Human MSC isolation and expansion

Human bone marrow-derived mesenchymal stromal cells (MSC) were obtained from the Center for Modular Manufacturing of Structural Tissues at Case Western Reserve University (Cleveland OH). The process used for isolating cells has been described previously [32,33]. Isolated cells were expanded in DMEM with low glucose (Gibco, Cat # 11885084) supplemented with 10 % fetal bovine serum (Gibco, Cat # 26140-079) and 10 ng/mL fibroblast growth factor 2 (Shenandoah Biotechnology, Part # 100-146). MSC used in subsequent experiments were at passage 6 (9-11 population doublings).

Primary human umbilical vein endothelial cells (EC; Lonza, Inc., Walkersville, MD) were expanded in fully supplemented endothelial cell growth medium (ECG; VascuLife® VEGF Endothelial Medium Kit, LifeLine Cell Technology). Cells were cultured in T-175 flasks to 80 % confluence and maintained at 37 °C, 5 % CO_2_, and 95 % humidity with medium changes every 48 hours.

### Generation of chondrogenic and hypertrophic MSC pellets

MSC culture flasks reached confluency after 6 days of expansion. At this point, cells were trypsinized and resuspended in chondrogenic differentiation medium (CDM,consisting of High Glucose DMEM (Gibco, Cat # 11965-092), supplemented with 1 % (v/v) insulin-transferrin-selenium (Corning, Cat # 354350), 1 % (v/v) non-essential amino acids (Gibco, Cat # 11-140-050) 40 *µ*g/mL L-proline (Sigma, cat # P-0380), 50 *µ*g/mL L-ascorbic acid-2-phosphate (Sigma, Cat # A8960), 0.1 *µ*M dexamethasone (Sigma, Cat # D4902), and 10 ng/mL TGF-*β* (Shenandoah Biotechnology, Part # 100-39). Pellets were formed by dispensing 2×10^6^ cells into individual wells (200 K cells/well) of a 96-well plate (Fisher, Cat # 08-772-54) and centrifuging at 1640 rpm for 5 min. Plates were then incubated at 37 °C, 5 % CO_2_, and 95 % humidity for 72 hours before first media change. Culture medium was changed every 48 hours for the remainder of culture.

The process employed to generate chondrogenic and hypertrophic pellets is outlined schematically in **Figure 1**. For the first 14 days of culture, pellets were primed in CDM. Pellets were then either maintained in CDM for an additional 7 days of culture to produce chondrogenic pellets or switched to hypertrophic differentiation medium (HDM) to generate hypertrophic pellets. HDM consisted of High Glucose DMEM, supplemented with 1 % (v/v) insulin-transferrin-selenium, 1 % (v/v) non-essential amino acids, 40 *µ*g/mL L-proline, 50 *µ*g/mL L-ascorbic acid-2- phosphate, 10 nM triiodothyronine, and 10 mM *β*-glycerophosphate.

**Figure 1:**
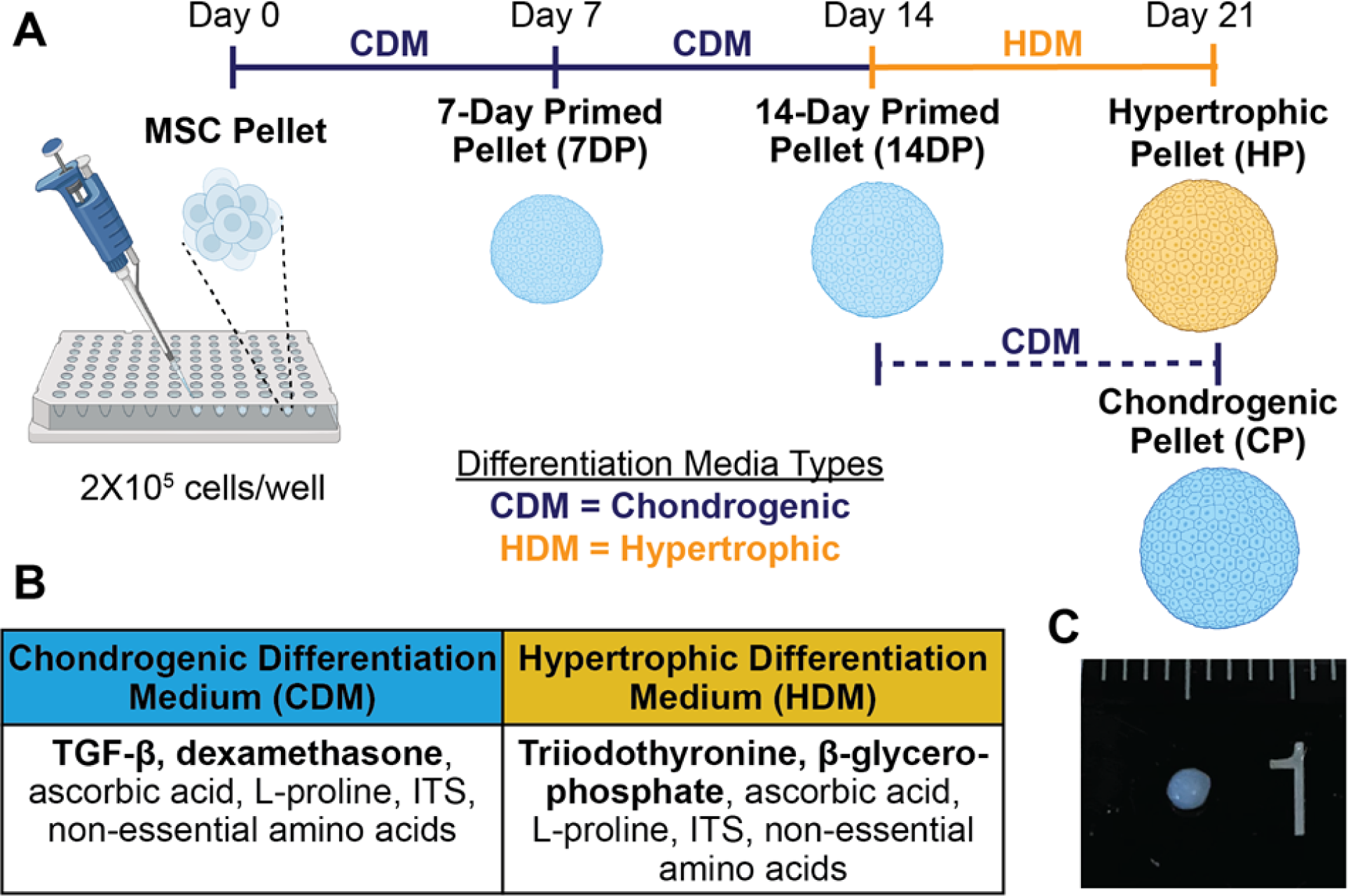
Generation of chondrogenic and hypertrophic hMSC pellets — A) Schematic illustration of the process used to generate chondrogenic and hypertrophic pellets (not to scale). B) Table of components for differentiation media used throughout the induction process. C) Representative image of HP.

### Quantification of glycosaminoglycan content

To assess the accumulation of cartilage-specific matrix in pellet cultures, levels of glycosaminoglycans (GAG) were quantified using the dimethylmethylene blue (DMMB) assay as described in previous work [34]. Briefly, pellet samples were homogenized and digested at 65 °C overnight in papain solution (125 *µ*g/mL papain, 50 mM sodium phosphate, pH 6.5). GAG content in the digested lysate of each sample was subsequently determined by comparing its DMMB reading to a standard curve. Assay was performed in triplicate.

### Histology analysis

Samples were fixed in 10 % aqueous buffered zinc formalin (Fisher), dehydrated in a graded series of ethanol solutions (70-100%), embedded in paraffin wax (Sigma-Aldrich), and sectioned at 7 *µ*m thickness. All stains were performed on middle sections of pellet cultures and imaged using an inverted microscope (Nikon Eclipse, Nikon USA).

To assess sulphated glycosaminoglycan (sGAG) content, sections were stained with 1.0 % (w/v) Alcian blue 8GX (Sigma) in 3.0 % acetic acid, with a counter stain of 0.1 % (w/v) nuclear fast red to assess cellular distribution. To visualize areas of matrix mineralization, slides were stained with 2 % alizarin red.

Immunofluorescent staining was performed to visualize specific matrix proteins and biologic markers. Sections were incubated with primary antibody against Collagen Type X (1:100, ABclonal, Cat # A6889), RUNX2 (1:100, ABclonal, Cat # A2851), MMP13 (1:200, ABclonal, Cat # A16920), and VEGF (1:100, ABclonal, Cat # A12303) overnight at 4 °C. Primary antibodies were diluted in 0.1 % TBS-TX100. Slides then stained with secondary antibody (1:500, Invitrogen, A11034) and 4’, 6-diamidino-2-phenylindol (DAPI, Sigma, 300 nM final) diluted in 0.1% TBS- TX100 for 1.5 h. All images were taken on an inverted microscope (Nikon Eclipse, Nikon USA).

### Growth factor secretion from pellet cultures

Conditioned media was collected from pellet cultures after 48 hours of culture and stored at −80 °C. These were screened for the presence of specific analytes (VEGF-A, Angiopoietin-2, and Endothelin-1) using a fluorometric MILLIPLEX ® Human Angiogenesis/Growth Factor Magnetic Bead Panel Multiplex Assay (Sigma-Aldrich, Cat # HAGP1MAG). Each plate was run according to the manufacturer’s protocol using 25 *µ*L sample per well.

### Vasculogenic culture of pellets

A vascular matrix was formed by co-encapsulating EC and MSC within a fibrin carrier (Sigma-Aldrich, 2.5 mg/mL final clottable protein) and casting the cell-laden construct into individual wells of a 48-well plate (250 *µ*L/well). Fabrication of the vascular matrix involved suspending EC and MSC (1:1, 5 × 10^5^ total cells/mL) in a mixture of fibrinogen (2.5 mg/mL final clottable protein), fetal bovine serum (FBS, 10% v/v final, Corning), and thrombin (1 U/mL final, Sigma). Fibrinogen stock solution was prepared by dissolving the lyophilized protein in PBS (4.0 mg/mL clottable protein) and passing through a sterile filter. To assess the influence of paracrine signals from pellet cultures, Transwell® inserts, containing either 7-day primed (7DP) pellets, hypertrophic pellets (HP) or no pellets (control), were placed on top of each construct. Preparation of the inserts involved placing 10 pellets of the respective condition within the insert (or no pellets for the control condition) and casting a thin fibrin gel to hold the pellets in place. The entire Transwell® - vascular matrix construct was then incubated for 30 min at 37 °C, 5 % CO_2_, and 95 % to facilitate complete gelation. Each construct received 0.5 mL of either ECG or basal medium (High glucose DMEM with 2 % FBS) for 14 days of culture and medium was changed every 48 hours throughout culture.

### RNA isolation and gene expression analysis

Pellets were harvested at specified time points throughout culture for gene expression analysis using qRT-PCR. For each sample, five pellets were pooled, rinsed with PBS, snap frozen in liquid nitrogen, and stored at −80 °C. For RNA extraction, samples were homogenized with TRIzol RT Reagent (Molecular Research Center, RT 111) and pulverized by mechanical compression. Bromoanisole was added for phase separation and isolation of RNA. Purified RNA was then suspended in ultra-pure DI water before being quantified spectroscopically (Nanodrop 2000c Spectrophotometer, Thermo Fisher). Reverse transcription was performed utilizing a High-Capacity cDNA Reverse Transcription Kit (Applied Biosystems, Cat # 4368814) according to the manufacturer’s protocol. Gene expression analysis was performed on Applied Biosystems real-time PCR system using SYBR Green Reaction Mix (Applied Biosystems, Cat # 4385612). Relative expressions of aggrecan (ACAN), matrix metalloproteinase-13 (MMP13), runt-related transcription factor 2 (RUNX2), collagen Type 10 (COL10), SRY-Box Transcription Factor 9 (SOX9), alkaline phosphatase (ALP), vascular endothelial cell growth factor (VEGF), osterix (OSX), osteocalcin (OCN), and Dentin Matrix Acidic Phosphoprotein 1 (DMP1) were calculated and analyzed using the ΔΔCT method. TATA-box binding protein (TBP) and beta-glucuronidase (GusB) were used as housekeeping genes. The sequences of primers for each gene are listed in **Table 1**.

**Table 1:**
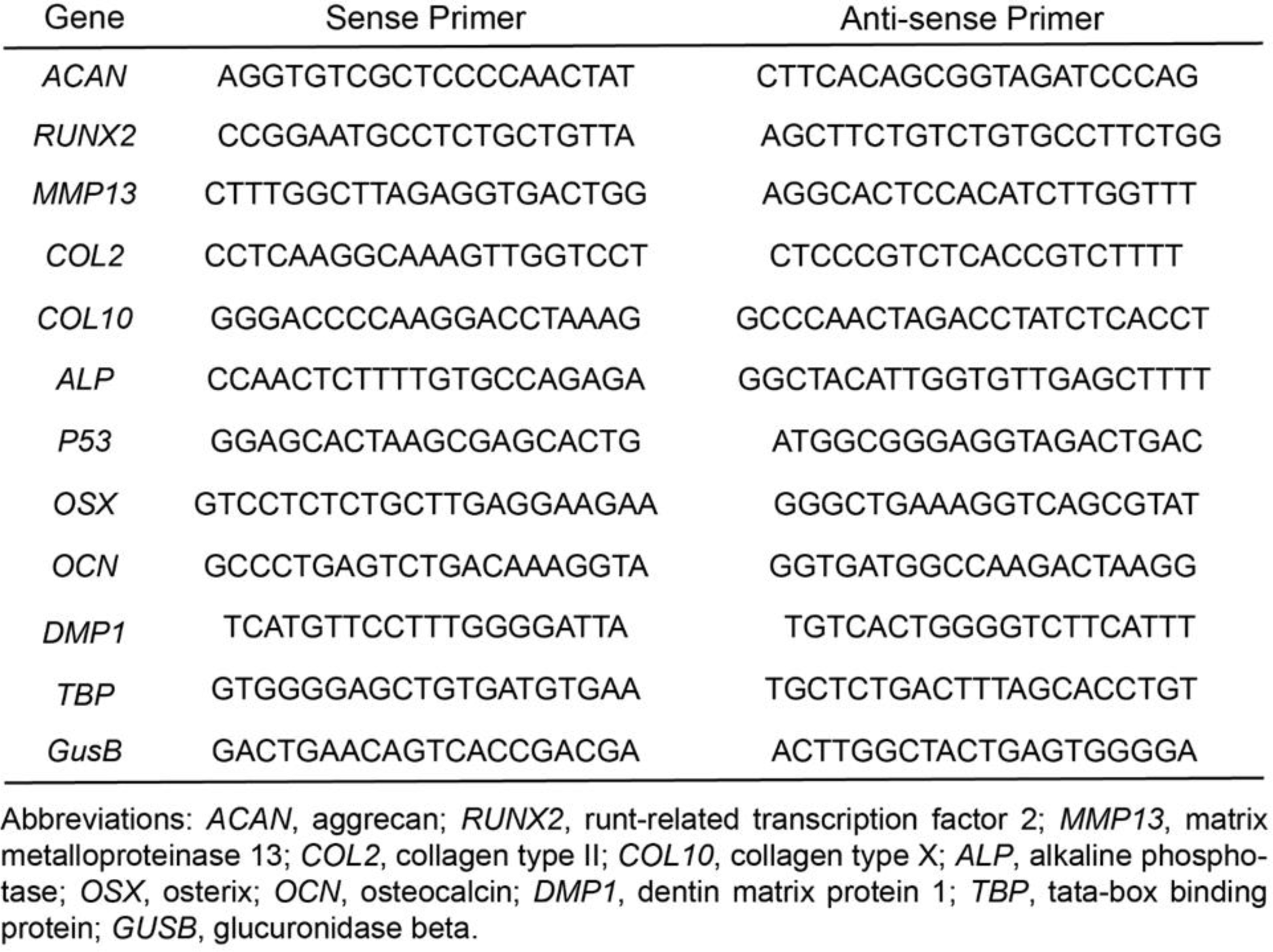
Primer sequences for qPCR gene expression analysis.

### Fabrication of Vasculogenic (VAS) Microtissues

Vasculogenic microtissues comprised of a fibrin hydrogel containing MSC and EC (1:1 cell ratio) were fabricated using a water-in-oil emulsification process, as previously described [11,15]. Briefly, fabrication involved suspending MSC and EC in a mixture of fibrinogen (Sigma-Aldrich), fetal bovine serum (FBS, Corning), and thrombin (50.0 U/mL, Sigma). Fibrinogen stock solution was prepared by dissolving the lyophilized protein in PBS (4.0 mg/mL clottable protein) and passing through a sterile filter. Microtissues were generated in batches of 6 mL. For each batch of microtissues, 1.53 mL of a cell suspension containing MSC and EC (1:1, 2 × 10^6^ total cells/mL), 600 *µ*L FBS (10% v/v final), 120 *µ*L thrombin (1 U/mL final), and 3.75 mL fibrinogen stock solution (2.5 mg/mL final clottable protein) were added and mixed thoroughly before emulsification. The mixture was then dispensed into a polydimethylsiloxane (PDMS) bath and mixed by a dual radial-blade impeller at 600 rpm for 5 min to allow for emulsion. Subsequently, the bath temperature was increased to 37 °C and the emulsion mixture was stirred for an additional 25 min to achieve full gelation. Density separation was used to isolate formed microtissues from the PDMS solution. Microtissues were then resuspended in ECG and transferred to vented conical tubes (CellTreat). Tubes were incubated at 37 °C, 5 % CO_2_, and 95 % for 5 days in ECG to preculture, with media changed every 48 hours.

### Fabrication of Multiphase Constructs

Multiphase constructs contained HP and VAS microtissues homogenously distributed throughout a bulk fibrin hydrogel carrier. Generation of constructs first involved placing 5 hypertrophic pellets within a single well of a 48-well plate. VAS microtissues were then suspended in a mixture of fibrinogen, FBS, and thrombin, and cast on top of HP (250 *µ*L/well). All stock solutions were kept on ice prior to construct fabrication. Fibrinogen stock solution was prepared by dissolving lyophilized protein in PBS (4.0 mg/mL clottable protein) and passing through a sterile filter. Batches of microtissues within the fibrin precursor mixture were made 2 mL at a time. For each batch, 510 *µ*L of microtissues (containing 2 × 10^6^ cells), 200 *µ*L FBS (10% v/v final), 40 *µ*L thrombin (1.0 U/mL final), and 1.25 mL fibrinogen stock solution (2.5 mg/mL final clottable protein) were combined and mixed thoroughly. For each batch, seven constructs (250 *µ*L each) were cast into each well. The plate was then incubated for 30 min at 37°C, 5 % CO_2_, and 95 % to facilitate complete gelation of the bulk carrier gel. Each construct received 0.5 mL of either endothelial cell growth (ECG) medium or basal medium (High glucose DMEM with 2 % FBS).

The method to achieve the desired cell and microtissue quantities per construct was modified from a previously published study [11]. Briefly, microtissues were re-suspended in 2.0 mL of DMEM and transferred into vented 15 mL conical tubes with filters (CELLTREAT). A 100 microliter sample of the microtissue suspension was transferred to a 96-well plate and incubated with 100 *µ*L of nattokinase (1.0 mg/mL in PBS containing 1.0 mM EDTA, Japan Bio Science Laboratory Co., Ltd) to degrade the microtissue matrix and dissociate the embedded cells. The plate was incubated for 30 min at 37 °C to facilitate enzyme activity and microtissue degradation. Cells suspended in the solubilized matrix were then counted using a hemocytometer and the cell count was used to calculate cell density per microtissue volume of the remaining batch. The final volume of the microtissue suspension was then adjusted to provide appropriate cell numbers per construct.

### Quantification of Vessel Formation

Constructs were rinsed with PBS 3X for 5 min each and fixed in aqueous buffered zinc formalin (Z-fix, Fisher) for 30 min at room temperature. After removal of fixative and rinsing in PBS, constructs were stained with of a staining solution comprised of Ulex Europaeus Lectin 1 (UEA, Vector Laboratories) diluted in PBS (1:100) and DAPI (1:500). The entire culture plate was wrapped in foil and incubated overnight at 4 °C. Gels were then rinsed with PBS at room temperature and stored at 4 °C until imaged.

Imaging was performed using an Olympus IX81 equipped with a Disc Spinning Unit and a 100 W high-pressure mercury burner (Olympus America, Center Valley, PA), a Hamamatsu Orca II CCD camera (Hamamatsu Photonics, K.I., Hamamatsu City, Japan), and Metamorph Premier software (Molecular Devices, Sunnyvale, CA). For vessel network formation, 5 images per construct were taken over a 200 *µ*m depth (10 slices at 20 *µ*m thick) at 4X magnification. Slices were superimposed using maximum intensity z-projection. Total network density was quantified using the Angiogenesis Tube Formation module in Metamorph Premier (Molecular Devices).

### Analysis of Osteogenic Differentiation

Intracellular alkaline phosphatase (ALP) activity, DNA, and calcium deposition were quantified to assess the osteogenic phenotype of cells. To assess osteogenic activity in pellets, two pellets were pooled per sample. Samples were rinsed with PBS before incubating in 0.5 % TBS-TX100 to lyse cellular membrane to generate the sample solution used in subsequent assays. To analyze osteogenic activity in multiphase constructs, samples were centrifuged at 1 × 10^4^ g for 5 min to remove excess media within the hydrogel. Samples were then rinsed with PBS before adding 500 μL of 0.5 % TBS-TX100 to lyse cellular membrane and generate sample solution used in subsequent assays. To facilitate digestion, both pellet and construct sample solutions were sonicated for 10 sec at 10% power. Sample solution was used for all subsequent ALP, DNA, and calcium assays.

Intracellular ALP activity was quantified using a commercially available assay (Sigma). Standard curves were constructed using serial dilutions of p-nitrophenol standard solution (Sigma, N7660) in a 2.0 N NaOH solution. The assay was performed in a clear 96-well plate. Triplicate sample wells each contained 200 μL of appropriate standard solution, 10 μL of sample solution and 100 μL working solution. The working solution was comprised of ALP substrate solution (Sigma, N2765) diluted with an equal part of Alkaline Buffer Solution (Sigma). After 30 min of incubation at 37 °C, 90 μL of 2.0 N NaOH was added to each sample-well to stop the reactions and absorbance was subsequently read at 405 nm.

DNA content was quantified using a standard DNA assay kit (Quant-iT™ PicoGreen dsDNA kit; Invitrogen) following the manufacturer’s protocol. The cell lysate solution was prepared in the same manner as for the ALP assay. Standard curves were constructed using serial dilutions of λDNA standard (Invitrogen, 25250) in a 10 mM Tris-HCl solution (pH 7.5). The assay was performed in a black 96-well plate. Triplicate sample wells each contained 100 μL of 10 mM Tris-HCl, 10 μL of sample and 110 μL of PicoGreen buffer. Buffer was made by 200-fold dilution of Quant-iT PicoGreen dsDNA Reagent (ThermoFisher, P11495) in 10 mM Tris-HCl. The plate was covered in foil and incubated for 30 min at 37 °C. Following incubation, the plate was read at an excitation of 498 nm and emission of 518 nm.

Calcium deposition was quantified using the orthocresolphthalein complex one (OCPC) assay as previously described [9]. Standard curves were constructed using serial dilutions of CaCl_2_ (Sigma) dissolved in 1.0 N acetic acid. The assay was performed using a clear 96-well plate. Triplicate sample wells each contained 20 μL of sample, and 250 μL of working solution consisting of 0.05 mg/mL OCPC in ethanolamine-boric acid−8-hydroxyquinoline buffer. After 10 min of incubation at room temperature, absorbance was measured at 575 nm.

### Statistical analysis

Data are presented as mean±SD (n=3). Statistical significance was determined using 2-way analysis of variance (ANOVA) with a Tukey’s multiple comparison post hoc test. All statistical analysis was performed using Prism 8.3 software (GraphPad, San Diego, CA). P<0.05 was considered statistically significant.

## 3. Results

### Induction of hypertrophy in MSC-derived chondrocytes

Scaffold free pellet culture was used to promote chondrogenic differentiation of hMSC. Phenotype characterization data are shown in **Figure 2**. DNA content and matrix accumulation were assessed throughout culture to characterize chondrogenic behavior and pellet formation. Pellets maintained in chondrogenic differentiation medium (CDM) for the entire 21-day culture period displayed no change in total DNA content, whereas those switched to hypertrophic differentiation medium (HDM) at day 14 exhibited a statistically significant decrease in DNA levels by Day 21 (Fig. 2A). Culture in CDM supported a significant increase in sulphated glycosaminoglycan (sGAG) accumulation by Day 14 (Fig. 2B). In the last week of culture, both pellets maintained in CDM and those switched to HDM exhibited a sustained increase in sGAG accumulation by Day 21, leading to the formation of chondrogenic pellets (CP) and hypertrophic pellets (HP), respectively. Hypertrophic induction led to less total sGAG content within pellets, both in absolute terms and when normalized to DNA levels (Fig. 2C). HP were found to be statistically smaller in size compared to CP, with average diameters of 1.55±0.13 and 1.72±0.07 mm, respectively (Fig. 2D). Alcian blue histological evaluation revealed that CP and HP were uniformly spherical in morphology with dense sGAG homogenously distributed throughout the pellet (Fig. 2E). Microscopic observation revealed hypertrophic-like morphology in HP based on the enlarged cells and lacunae compared to CP.

**Figure 2:**
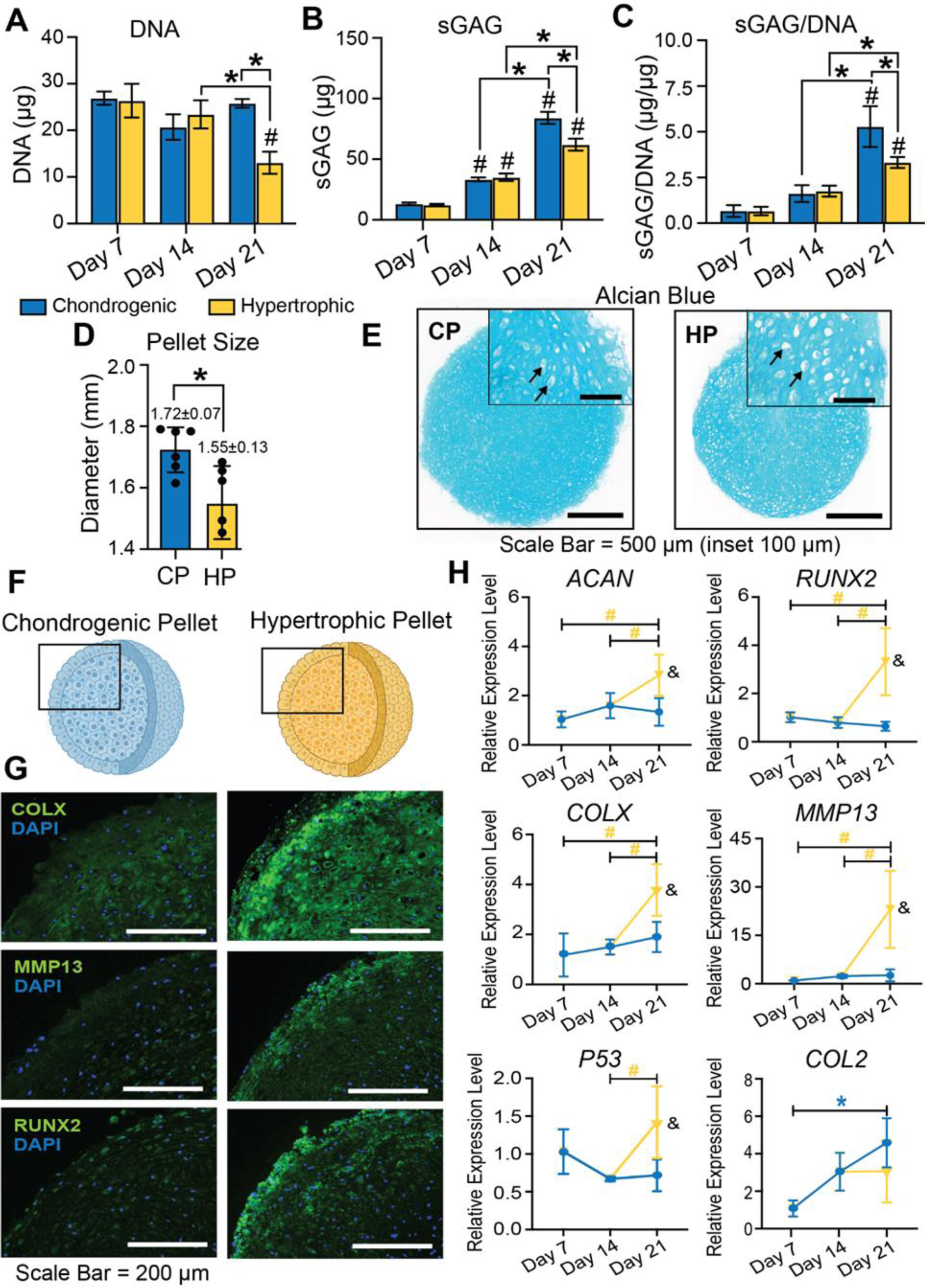
Characterization of hypertrophic pellets — Levels of A) DNA, B) sGAG, and C) DNA-normalized sGAG were quantified at specific time points throughout the culture period. # indicates statistical significance compared to day 7 of the same condition. * indicates statistical significance between specified groups. D) The average diameter of pellets with and without hypertrophic induction measured at day 21. E) Alcian blue staining at day 21 of culture. Black arrows denote lacunae. F) Schematic illustration of pellets showing regions imaged for histological analysis as shown in black box (not to scale). G) Immunofluorescent staining of COLX, MMP 13, and RUNX2. Blue = nuclei. H) Quantitative analysis of chondrogenic and hypertrophic genes, ACAN, RUNX2, COLX, MMP13, P53, and COL2. # indicates statistical significance of HP between specified groups. & indicates statistical significance between HP and CP of the same time point.

Pellets were then examined for markers of hypertrophy throughout the culture period to assess phenotype progression. Immunofluorescence analysis for the hypertrophic markers collagen type-X (COLX), matrix metalloproteinase-13 (MMP13), and runt-related transcription factor 2 (RUNX2) revealed high expression of these markers in HP whereas CP exhibited very little to no expression, indicating onset of hypertrophy in HP (Fig. 2F&G). Gene expression analysis revealed an upregulation of the chondrogenic marker *ACAN* and the hypertrophic markers *RUNX2*, *COL10*, *MMP13*, and *P53* in HP compared to CP through the 7-day induction period (Fig. 2H). No change in expression levels of *COL2* was observed in HDM, though pellets that were maintained in CDM displayed a significant increase by day 21. Taken together, this data suggests successful progression of HP towards endochondral ossification pathway.

### Hypertrophic pellets exhibit osteogenic activity

The capacity of hypertrophically-induced pellets to support osteogenic activity and mineralization by hMSC within HP was investigated, as shown in **Figure 3**. Alkaline phosphatase (ALP) activity, an early-stage marker of osteogenic differentiation, exhibited a significant increase from day 14 to day 21 in culture, whereas ALP activity in CP decreased over the same time course (Fig. 3A). Calcium deposition was quantified to assess pellet mineralization, and only HP exhibited a statistical increase in mineral levels by day 21 (Fig. 3B). This pattern was confirmed through alizarin red histological analysis, which showed robust mineral deposition in HP and no observable positive staining in CP (Fig. 3C). Mineral deposition was most prominent near the periphery of HP, with nodule formation also being observed sparsely throughout the center of the construct. Quantiative gene expression analysis revealed upregulation of osteogenic genes *ALP*, *OSX*, *OCN*, and *DMP1* in HP by day 21 of culture, whereas pellets maintained in CDM displayed no statistical change in expression throughout the culture period (Fig. 3D). These results confirm the ability of HP to mineralize the accumulated cartilage ECM, while also providing evidence of possible phenotype plasticity of HP towards an osteogenic lineage.

**Figure 3:**
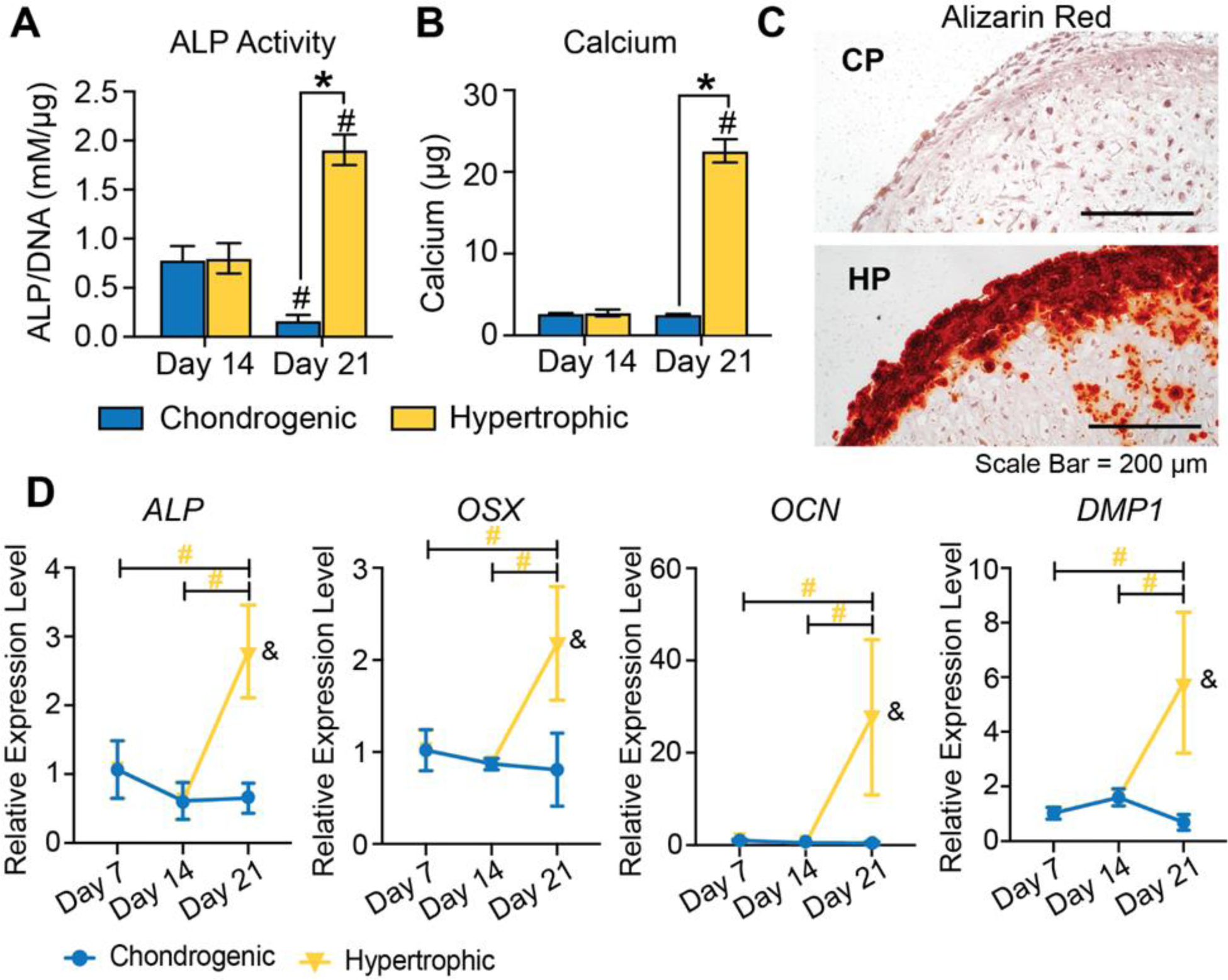
Effect of hypertrophic induction on osteogenic activity — Quantification of A) ALP activity and B) calcium deposition within constructs. # indicates statistical significance compared to day 14 of the same condition. * indicates statistical significance between specified groups. C) Alizarin Red staining of hypertrophic pellets (HP) and chondrogenic pellets (CP) at day 21. D) ALP, OSX, OCN, and P53 gene expression. # indicates statistical significance of HP between specified groups. & indicates statistical significance between HP and CP of the same time point.

### MSC pellets produce and secrete angiogenic factors

The secretion of specific angiogenic factors known to promote vessel development during endochondral bone formation was examined, as shown in **Figure 4**. The presence of angiotensin- 2, endothelin-1, and VEGF-A in pellet-conditioned medium was analyzed at defined time points to interpret relative degrees of secretion throughout the culture period. Pellet formation and chondrogenic priming supported the production and secretion of all tested factors (Fig. 4A). In particular, hypertrophic induction enhanced the secretion of angiotensin-2 and endothelin-1 by HP while secretion levels of CP remained unchanged by day 21. Interestingly, high levels of VEGF-A were present at 7 days of chondrogenic priming, but were found to significantly decrease by day 14, even in HP. A key characteristic of native endochondral ossification is the sequestration of VEGF and other factors in the matrix to create gradients that guide vessel infiltration. Qualitative analysis of immunofluorescence (Fig. 4B) showed increased sequestration of VEGF-A over time in culture (Fig. 4A), suggesting that the released growth factor was bound by the pellet matrix, especially in HP. These results demonstrate that hypertrophic induction increases the secretion and sequestration of angiogenic factors by MSC pellets, which may support vessel development.

**Figure 4:**
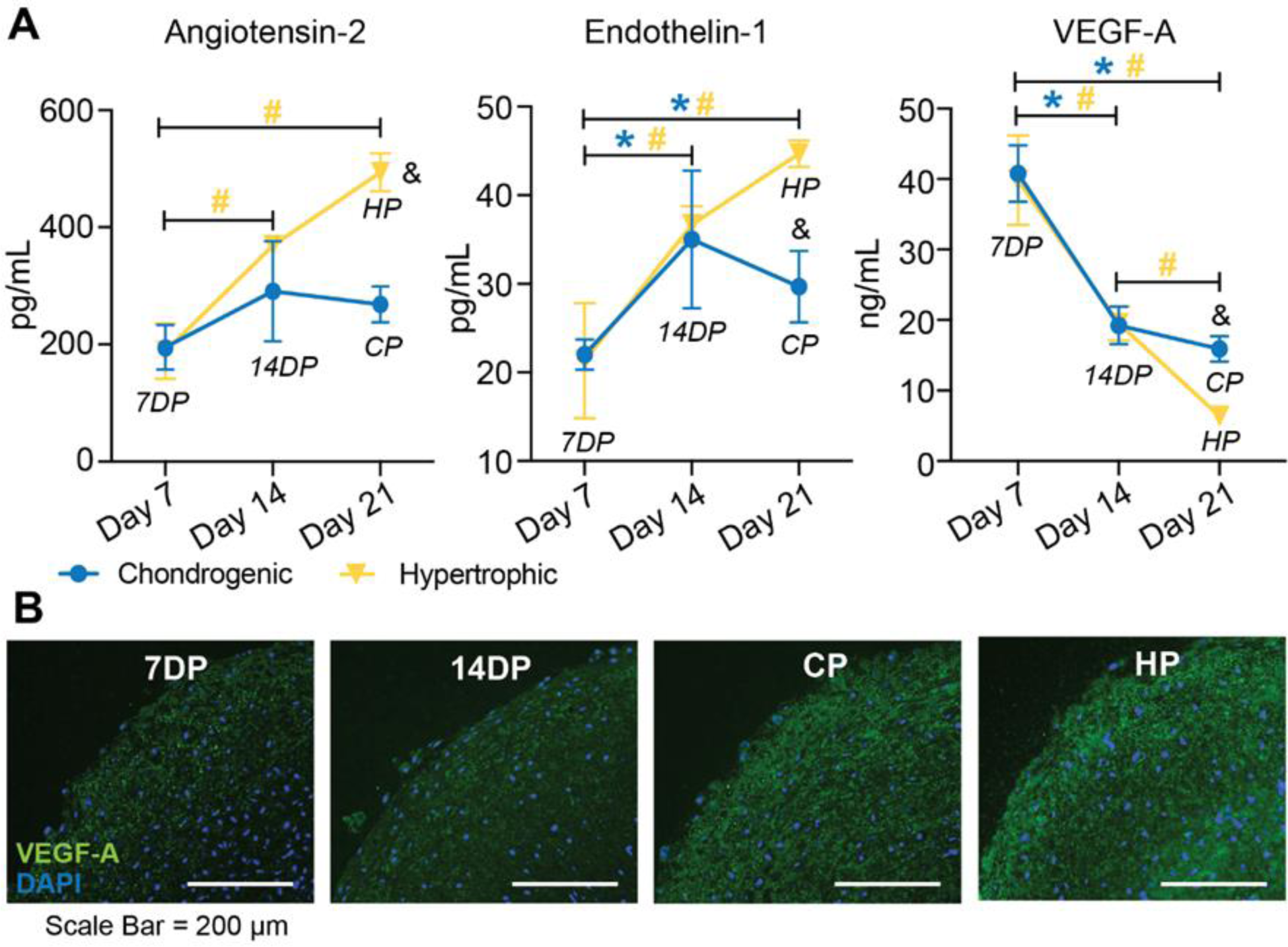
Angiogenic factor secretion throughout the culture period — A) Quantification of angiotensin-2, endothelin-1, and VEGF-A secretion from pellets. 7DP = 7-day primed pellets; 14DP = 14-day primed pellets; CP = Chondrogenic pellets; HP = Hypertrophic pellets. * indicates statistical significance between chondrogenic pellets of specified time points. # indicates statistical significance between hypertrophic pellets of specified time points. & indicates statistical significance between chondrogenic and hypertrophic pellets at the specified time point. B) Immunofluorescent staining for VEGF-A in constructs throughout the culture period.

### Hypertrophic pellets support new vessel development

The capacity of hypertrophic chondrocytes to support the formation of new vessel-like structures was examined in a co-culture system, as shown in **Figure 5**. Either 7-day primed (7DP) pellets or HP were cultured in a transwell insert in a surrounding fibrin matrix seeded with MSC and EC. In this system, the pellets and vascularizing cells were separated by a semipermeable membrane to permit exchange of secreted factors (Fig. 5A). The entire system was cultured in either endothelial cell growth (ECG) or basal medium for 14 days and subsequently imaged using confocal microscopy to evaluate vessel development. All constructs cultured in ECG medium formed vessel-like structures, regardless of pellet condition (Fig. 5B). When cultured in basal medium that lacked angiogenic factors, only constructs containing pellets displayed similar vessel-like structures. Quantification of vessel network density revealed that although constructs with HP cultured in ECG medium contained statistically fewer vessels compared to those with no pellets, both 7DP pellets and HP cultured in basal medium exhibited statistically more vessels compared to those containing no pellets (Fig. 5C). These data demonstrate the intrinsic capability of HP to support vessel formation without requiring addition of exogenous angiogenic growth factors.

**Figure 5:**
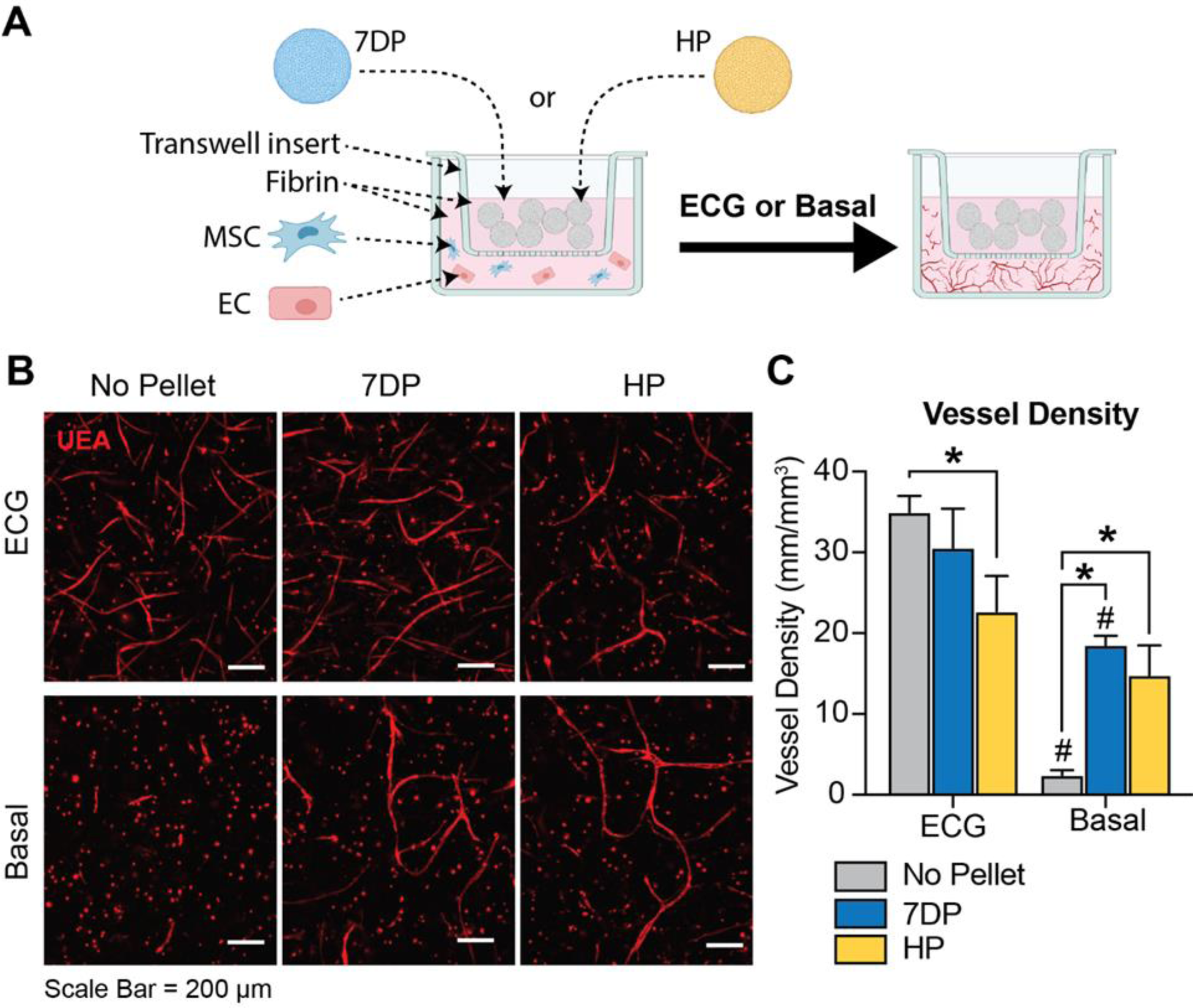
Support of vessel formation by pellets in a co-culture system — A) Schematic illustration of experimental setup. 7DP or HP were cultured with a vascular matrix containing MSC and EC in either endothelial cell growth (ECG) or basal medium for 14 days. The pellets separated from the vascular matrix by semipermeable membrane (Transwell® insert). B) Confocal microscopy images of constructs at day 14 stained with UEA (red), revealing formed vessel-like structures. C) Quantification of vessel density. # indicates statistical significance compared to ECG group of the same condition. * indicates statistical significance between specified groups.

### Multiphase constructs support vessel development and osteogenic activity

Multiphase constructs were fabricated to examine the ability for HP to support concomitant development of both vascularized and mineralized engineered tissue without addition of exogenous angiogenic growth factors or inductive components, as shown in **Figure 6**. Multiphase constructs were generated by co-encapsulating HP and vascular microtissues (V*µ*T) within a surrounding fibrin gel and were cultured for 14 days in either ECG or basal medium (Fig. 6A). Constructs were imaged at the end of co-culture using confocal microscopy to visualize vessel formation (Fig. 6B), revealing varying degrees of vessel development based on the presence of HP and type of culture medium (Fig. 6C). As expected, constructs containing only V*µ*T displayed extensive EC sprouting when cultured in ECG medium, whereas negligible vessel-like structure development was observed when cultured in basal medium. While multiphase constructs with HP also supported EC sprouting from V*µ*T when cultured in ECG medium, those cultured in basal medium displayed a more extensive network of vessel-like structures compared to constructs without HP. Quantification of network density within constructs confirmed the qualitative observations (Fig. 6D). Constructs containing only V*µ*T contained statistically fewer vessels when cultured in basal medium as compared to those that were cultured in ECG medium. In ECG medium, the addition of mineralizing HP to create multiphase resulted in decreased vessel formation relative to constructs containing only V*µ*T. However, multiphase constructs containing HP were found to significantly enhance vessel development when cultured in basal medium, further confirming the intrinsic angiogenic capabilities of HP. Multiphase HP-containing constructs cultured in basal medium had similar vessel density to V*µ*T-only constructs cultured in ECG, which displayed the highest degree of vessel development out of all tested conditions.

**Figure 6:**
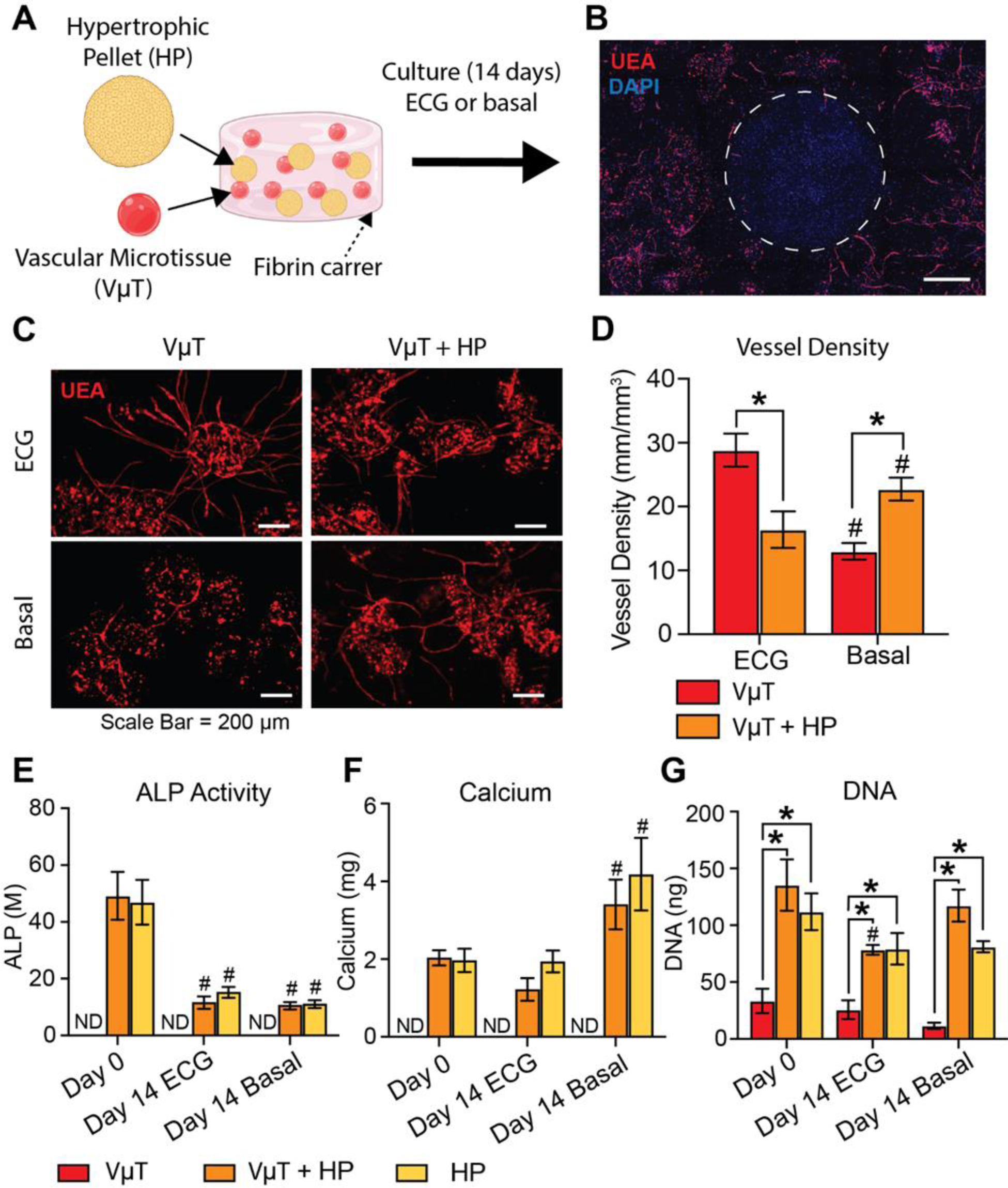
Osteogenic and vasculogenic activity of multiphase constructs composed of hypertrophic pellets (HP) and vascular microtissues (VμT) — A) Schematic illustration showing fabrication and culture process used to generate multiphase constructs. HP were encapsulated with vascular microtissues within a fibrin gel and cultured in ECG or basal medium for up to 14 days. B) Representative image of multiphase construct at day 14, depicting formed vessels (UEA, red), nuclei (DAPI, blue), and HP (outlined with dotted circle). C) Confocal images of tested conditions at day 14, depicting formed vessels (red). D) Vessel density quantification. # indicates statistical significance compared to ECG group of the same condition. * indicates statistical significance between specified groups. Quantification of E) ALP activity, (F) calcium deposition, and G) DNA content. ND indicates no detectable levels. # indicates statistical significance compared to day 0 of the same condition. * indicates statistical significance between specified groups.

The same construct conditions were then analyzed for markers of osteogenic activity at days 0 and 14 of co-culture. Multiphase constructs displayed a statistical decrease in ALP activity when cultured in either ECG or basal media (Fig. 6E). Both multiphase and HP-only constructs displayed a significant increase in calcium deposition when cultured in basal medium while those cultured in ECG remained unaffected (Fig. 6F). The addition of V*µ*T in multiphase constructs did not appear to influence either ALP activity or mineralization. As expected, constructs containing V*µ*T- only did not display any detectable levels of ALP or mineral deposition throughout culture. Multiphase constructs displayed a statistical decrease in total DNA content when cultured in ECG, though levels were maintained when cultured in basal media (Fig. 6G). Overall, the data indicates that HP in basal medium culture can support matrix mineralization of multiphase constructs without need for exogenous factors or mineral supplements.

To more completely characterize the capacity of the multiphase HP-containing system to support endochondral tissue development, gene expression in hypertrophic pellets was examined over the 14-day culture period, as shown in **Figure 7**. Hypertrophic cells cultured in basal medium maintained levels of *RUNX2* expression over time, whereas those cultured in ECG displayed decreased expression by day 14 in culture. *COL10* expression appeared decreased over time in culture regardless of which media type was used. Interestingly, HP-containing constructs culture in basal medium supported a significant (>10-fold) increase in *MMP13* and *OCN* expression levels by day 14. *ALP* gene expression was significantly decreased in both media conditions, displaying a similar trend to ALP activity. Expression of the bone transcription factor *OSX*, tumor protein *P53*, and extracellular matrix protein *DMP1*, were all downregulated over time in ECG medium, while expression levels of these genes in basal medium stayed constant over 14 days. Taken together, these data demonstrate the detrimental effects of ECG medium on the ability of HP to sustain concomitant osteogenic and as angiogenic capabilities. In addition, they provide support for the concept of harnessing the intrinsic angiogenic and mineralization abilities of HP-containing constructs to create vascularized engineered bone.

**Figure 7:**
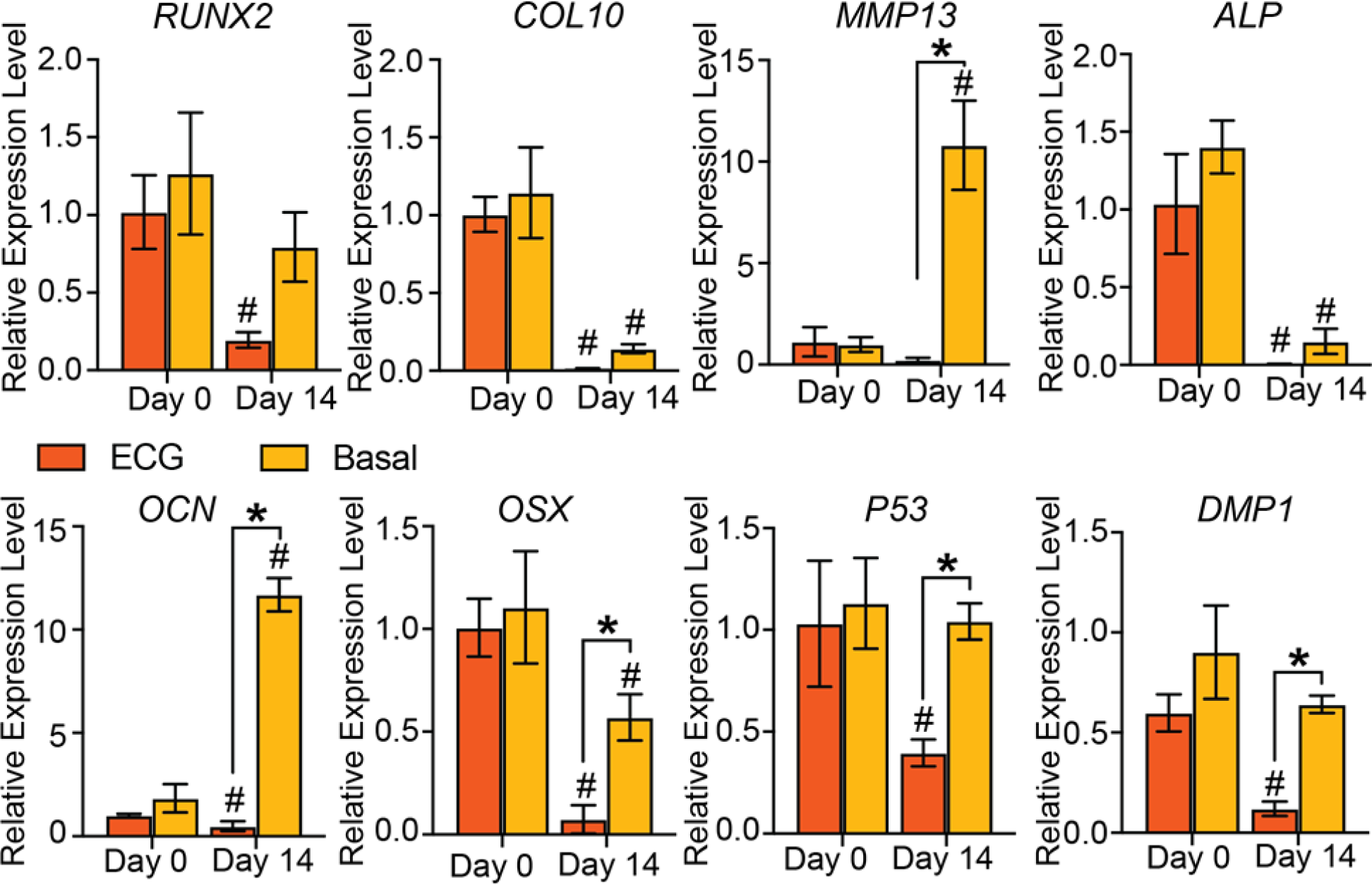
Expression of chondrogenic and osteogenic genes by HP in multiphase constructs — Expression levels of select genes in HP within the multiphase system analyzed throughout culture in either ECG or basal medium. # indicates statistical significance compared to day 0 of the same media condition. * indicates statistical significance between specified groups.

## 4. Discussion

This study characterized the development of a new approach to generating a vascularized bone construct *in vitro* by employing a modular biomaterials-based strategy and leveraging the regenerative phenotype of hypertrophic MSC-derived chondrocytes. Chondrogenic pellet culture of MSC and subsequent induction of hypertrophy generated modular units that demonstrated matrix remodeling, mineralization, and angiogenic behavior that mimicked native endochondral development. Modular vasculogenic microtissues were also created by combining a co-culture of MSC and EC within discrete fibrin microtissues, which demonstrated pericyte-like lineage commitment and robust vascular development in previous studies [35]. Hypertrophic pellets (HP) were combined with vasculogenic microtissues (V*µ*T) to generate multiphase constructs that combined both osteogenic and vasculogenic components. Subsequent culture of multiphase constructs revealed that the inclusion of hypertrophic pellets supported concomitant vessel development and tissue mineralization. Notably, this tissue development occurred in basal culture medium that lacked exogenous growth factors and inductive components. In contrast, culture in endothelial growth medium negatively influenced the hypertrophic phenotype of MSC and led to impaired vessel development and matrix mineralization.

In previous work by our group, an alternative strategy that mimicked the process of intramembranous ossification was applied to creating multiphase vascularized bone constructs [35]. In this previous approach, osteogenic microtissues containing MSC were directly induced toward the osteogenic phenotype prior to combination with vascularizing microtissues. It was demonstrated that vessel development in multiphase constructs was supported in endothelial cell growth medium. However, the osteogenic phase did not promote vessel development, and sustained mineralization of the construct was inhibited by endothelial cell growth medium. Other groups have reported similar challenges in tailoring the culture environment of engineered bone constructs to support vessel development [6,7]. Hypertrophic induction of MSC-derived chondrocytes has therefore been proposed as a strategy to emulate the natural mechanisms and sequential differentiation process of native endochondral ossification (EO), which supports both tissue vascularization and ossification [17]. This strategy was combined with a modular tissue engineering approach in the current study, in an effort to emulate the native progression of endochondral bone formation.

It has been demonstrated previously that hypertrophy can be induced in chondrogenic pellet culture *in vitro,* and that such constructs recapitulate key morphogenetic steps of native EO when transplanted *in vivo* [36–38]. In the current study, MSC-derived chondrocyte pellets were induced to hypertrophy, and subsequent characterization revealed positive expression of hypertrophic markers and genes. Collagen Type X, a matrix component produced by hypertrophic chondrocytes, was present ubiquitously throughout formed pellets by the end of the induction process, whereas markers associated with cell-mediated mineralization were localized towards the periphery. These findings are comparable to other work using a similar induction process, which observed regions in which cells exhibited a hypertrophic phenotype as well as areas containing undifferentiated cells [37]. It should be noted that a similar pattern of differentiation exists during developmental regulation of the growth plate, in which a border zone of hypertrophic chondrocytes instruct surrounding mesenchymal cells to differentiate into osteoblasts, resulting in the formation of a bony collar [39]. The expression of hypertrophy-specific markers within formed pellets confirmed the adoption of a hypertrophic phenotype in pellets. However, markers of osteogenic differentiation, such as OCN, OSX, ALP, and mineral deposition, are also expressed in hypertrophic chondrocytes. To more specifically characterize osteogenic differentiation within formed pellets, the expression of dentin matrix protein 1 (*DMP1*), an important signaling molecule in the induction of osteoblast differentiation and bone formation [40], was measured. While it remains unclear if cells within hypertrophic pellets take on an osteogenic phenotype in the current study, there is an emerging role of chondrocyte-to-osteoblast transdifferentiation in endochondral bone formation suggesting a transient behavior of MSC phenotype in this setting [28].

As chondrocytes mature to hypertrophy during native EO, they produce an angiogenic secretome that facilitates vascular infiltration within the formed cartilaginous matrix. This process is of interest in the engineering of vascularized bone because it suggests hypertrophic cells as a source of angiogenic cues to enable *in vitro* vessel development, while obviating the need for exogenous growth factors. Characterization of the production of select angiogenic factors in hypertrophic pellets showed that vascular endothelial growth factor A (VEGF-A) was secreted into the surrounding environment throughout the induction process, and was sequestered within the surrounding extracellular matrix over time. VEGF is a potent angiogenic signal that couples hypertrophic cartilage remodeling, ossification, and angiogenesis during endochondral bone formation [41], and VEGF is known to bind to the cartilage extracellular matrix during native EO, such that it can be released through select matrix metalloproteinase activity to promote vascular infiltration [41]. The hypertrophic induction process was also found to promote secretion of angiotensin-2 and endothelin-1 in pellets. These factors stimulate neovascularization and can act directly on EC to modulate proliferation, migration and morphogenesis [42,43].

Pellets in co-culture with a vasculogenic matrix containing MSC and EC produced robust vessel development when cultured in basal medium that did not contain endothelial cell growth factors. VEGF secretion is likely a driving influence in the observed vasculogenic activity; however, the degree of vessel development produced by 7-day primed pellets was similar to that from hypertrophic pellets, which exhibited significantly reduced VEGF secretion. It is therefore likely that the broader milieu of angiogenic signals produced through hypertrophic induction plays a role in *in vitro* vessel development. Interestingly, hypertrophic pellets demonstrated significantly reduced vessel formation when cultured in endothelial cell growth medium, compared to basal medium. Previous work has suggested that a vasculogenic culture environment can impair the chondrocyte-to-bone transition [31], and therefore it may have affected the secretome of hypertrophic pellets. Furthermore, a global transcriptome analysis of secreted factors from engineered cartilage derived from MSC found two factors that elicited anti-angiogenic effects, Serpin E1 and Indian Hedgehog [44]. In addition to VEGF, endothelial cell growth medium contains a variety of potent growth factors, including epidermal growth factor and fibroblast growth factor, both of which have been shown in previous to impair chondrocyte differentiation and maturation [45,46]. These results suggest that endothelial cell growth medium may promote the production of anti-angiogenic factors from hypertrophic pellets, and therefore that culture in basal medium provides an environment that is more conducive to hypertrophy-associated blood vessel development. This is particularly insightful as the use of endothelial growth medium is a widely used approach to maintain EC viability *in vitro* for the generation of a prevascularized bone construct [9,31,47], however this study highlights the deleterious effects of this method while providing an alternative by leveraging the vasculogenic paracrine signaling of hypertrophic- induced MSC without exogenous growth factors.

In the final stage of the present study, hypertrophic pellets were co-encapsulated with vascularizing microtissues to form a multiphase construct designed to support concomitant vascular and osseous tissue development. The guiding rationale in producing this multifunctional construct was that the hypertrophic cartilage component would secrete factors that support the formation of blood vessels by the vasculogenic component, while also undergoing mineralization and osteogenic differentiation, in a process that mimics endochondral ossification *in vivo*. The combined constructs demonstrated the ability to maintain cell viability and to support both robust vessel development as well as matrix mineralization throughout the *in vitro* culture period. Importantly, these effects were most evident when the constructs were maintained in basal culture medium, while the use of endothelial cell growth medium resulted in a decrease in cell number, fewer vessels and no change in mineral content. Furthermore, expression of genes associated with terminal hypertrophic differentiation and osteogenesis, P53 and DMP1, were maintained in basal medium culture, whereas constructs in endothelial growth medium displayed a decrease in expression. Similarly, culture of multiphase constructs in basal medium significantly increased expression levels of MMP13 and OCN, genes associated with matrix remodeling and mineralization, which may have promoted turnover of the formed cartilaginous and production of proteins that facilitate mineral deposition. Taken together, these results demonstrate the utility of a modular approach to generating multiphase engineered bone tissues, and highlight the ability to harness the process of endochondral ossification to enable the concomitant development of vascularized and mineralized tissues. This approach may alleviate some of the issues associated with the transplantation of osteogenic progenitor cells to achieve direct bone formation, and therefore has potential as an alternative cell-based therapy for regeneration of large and challenging bone defects.

## Acknowledgements

The authors gratefully acknowledge the assistance and guidance of their colleagues at the University of Michigan, and in particular Drs. Tiana Wong, Ciara Davis, Kurt Hankenson, Yadav Wagley, and Andrew Putnam. The authors also acknowledge the experimental support provided by the Center for Modular Manufacturing of Structural Tissue (P41 EB021911) at Case Western Reserve University, and particularly Dr. Rodrigo Somoza. This work was supported in part by the National Institute of Arthritis and Musculoskeletal and Skin Diseases (R01 AR062636, to JPS). NGS was partially supported by the Cellular Biotechnology Training Program (T32 GM008353) at the University of Michigan. The content is solely the responsibility of the authors and does not necessarily represent the official views of the National Institutes of Health. Some of the schematics in this paper were created using BioRender.com.

## Data Availability Statement

The data that support the findings of this study are available on request from the corresponding author,

